# Altair-dvOPM: an open-access platform for large-field three-dimensional tissue imaging

**DOI:** 10.64898/2026.05.08.723912

**Authors:** Tai Ngo, Muaaz Faiyazuddin, Trung Duc Nguyen, John Haug, Qionghua Shen, Seweryn Gałecki, Hazel M. Borges, Bingying Chen, Xiaoding Wang, Hao Zhu, Samuel S. Pappas, Fabian F. Voigt, Reto Fiolka, Kevin M. Dean

## Abstract

Altair-dvOPM is an open-access direct-view oblique plane microscope designed for large-field, three-dimensional imaging of cleared and expanded tissue sections. By combining photographic-lens-based detection, externally launched oblique illumination and precision-registered modular baseplates, the system achieves micrometer-scale lateral resolution over a ~5.4 mm field of view without custom objectives or highly specialized alignment procedures. We demonstrate imaging across scales, from subcellular structures in expanded cells to centimeter-scale expanded tissue sections, and provide documentation, CAD files, Zemax models and open-source control software to support replication and extension.

---

Chemically cleared and expanded cells and tissues are increasingly important for studies of tissue architecture, morphogenesis and cellular organization. However, their large size makes them challenging targets for high-resolution, three-dimensional fluorescence microscopy, often requiring long acquisition times to capture substantial volumes. Light-sheet fluorescence microscopy (LSFM) helps address these challenges by enabling gentle, parallelized, camera-based volumetric imaging, but its orthogonal illumination–detection geometry introduces additional complexity and limits compatibility with standard sample-mounting formats.

Single-objective approaches, including oblique plane microscopy^1^(OPM) and swept variants such as SCAPE^2^, address these limitations by using a single specimen-facing objective to collect fluorescence from an obliquely illuminated plane (Fig. 1A). The resulting oblique image plane is reimaged onto a camera through a remote-refocusing train, often implemented with secondary and tertiary objectives, enabling epi-style access and compatibility with conventional sample preparations while preserving the optical sectioning and volumetric throughput of LSFM. Together, these attributes have given rise to a diverse family of oblique-plane microscope architectures (see Supplementary Note 1)^3–9^.

**Figure 1:**
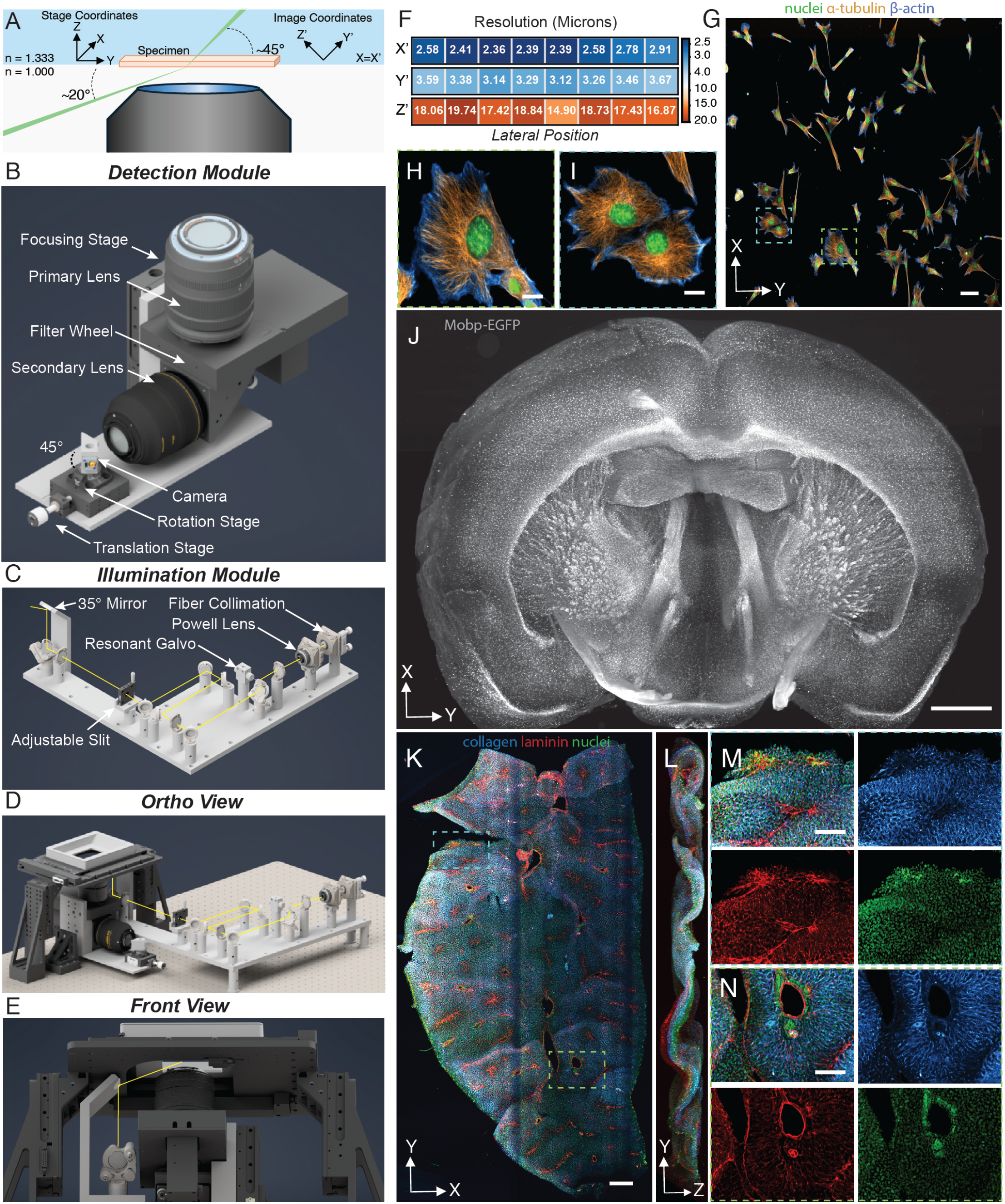
Altair-dvOPM design, resolution characterization and representative volumetric imaging performance. **A**, Schematic of the optical geometry. The illumination light-sheet is launched externally through the coverslip at an angle of approximately 20° and refracts to approximately 45° upon entering the aqueous specimen. Fluorescence from the obliquely illuminated plane is collected in the epi-direction by the photographic-lens-based detection path. Unprimed axes denote the stage coordinate system, in which *Y* is the sample-scanning direction, *Z* is the specimen-depth direction and *X* is parallel to the long axis of the light-sheet. Primed axes denote the oblique image coordinate system recorded by the camera, in which *Y*′ is the light-sheet propagation direction, *X*′ is the long axis of the light-sheet and *Z*′ defines the light-sheet thickness. The stage *X* axis corresponds to the image *X*′ axis. Thus, each camera frame captures an *X*′*Y*′ oblique plane, while volumetric imaging is performed by translating the specimen along stage *Y*. Following acquisition, image stacks are computationally sheared and rotated into the stage coordinate frame for visualization. **B**, Computer-aided design of the detection module, consisting of a monolithic baseplate mounted to a linear focusing stage, primary and secondary fast photographic lenses, an emission filter wheel and a rotated camera mounted on translation and rotation stages. **C**, Computer-aided design of the illumination module, consisting of fiber collimation optics, a Powell lens, a resonant galvanometer for shadow reduction and an adjustable slit for controlling the illumination numerical aperture. After light-sheet formation, the beam is folded vertically and then directed toward the specimen with a 35° mirror. **D**,**E**, Orthographic (**D**) and front (**E**) views of the assembled microscope. The sample is positioned on a stage platform that enables vertical specimen positioning and lateral sample scanning over a 120 × 75 mm travel range. **F**, Median full width at half-maximum (FWHM) values measured from images of 1 *µ*m fluorescent beads (*n* = 302 beads), binned by lateral position across the field of view. Values are reported in micrometers for the *X*′, *Y*′ and *Z*′ directions. **G**, Representative volumetric image of 4.2× expanded mouse embryonic fibroblasts stained for microtubules, *β*-actin and DNA. **H**,**I**, Zoomed in views of the regions indicated in **G. J**, Volumetric image of a chemically cleared sagittal brain section from a Mobp-eGFP mouse, with oligodendrocyte eGFP signal amplified by anti-GFP immunolabeling. **K**, Volumetric image of an entire 4× expanded mouse liver section stained for collagen I, laminin, and nuclei. **L**, Orthogonal projection of the image volume in **K. M**,**N**, Zoomed in views of the regions indicated in **K**. Scale bars, **G**, 100 *µ*m after expansion and 25 *µ*m before expansion; **J**, 1,000 *µ*m; **K**, 1,000 *µ*m after expansion and 250 *µ*m before expansion; **M** and **N**, 500 *µ*m after expansion and 125 *µ*m before expansion.

Conventional OPM relies on reimaging an oblique intermediate image plane with a tertiary imaging system, which is difficult to scale to large fields of view. Mesoscopic imaging requires low-NA reimaging optics because objective field of view generally increases as numerical aperture decreases; however, in traditional OPM geometries, fluorescence from the oblique image plane is efficiently relayed only when the reimaging NA exceeds approximately 0.5^9,10^. Below this limit, the obliquely propagating fluorescence falls outside the acceptance cone of the tertiary objective, leading to severe or complete signal loss^9,10^. These optical constraints are compounded by alignment sensitivity, as imaging performance depends on accurate pupil mapping between the specimen-facing and secondary objectives and precise alignment of the secondary and tertiary objectives. A recent direct-view variant avoided the tertiary-objective bottleneck by placing the camera sensor directly in the remote space, achieving a 5.3 × 2.6 mm field of view with ~2 *µ*m lateral and ~22 *µ*m axial resolution^11^. However, this implementation still requires a pupil-mapped relay to satisfy the constraints of remote refocusing.

An alternative approach is to use fast photographic lenses for remote focusing, which combine large fields of view, moderate numerical aperture and externally accessible pupils, providing a potential route to simplify oblique-plane imaging. Importantly, any OPM implementation must satisfy the constraints of remote refocusing to achieve aberration-minimized imaging, which requires that the lateral magnification between the specimen and remote spaces approximate the refractive-index ratio between the two media. We therefore surveyed commercially available photographic lenses and identified the Mitakon Speedmaster 65 mm f/1.4 and Nikon 85 mm f/1.4 lenses as a mechanically compatible pair with external pupil access, millimeter-scale imaging fields, a theoretical effective numerical aperture of 0.25 and a measured magnification of 1.307×, close to the magnification of *n*_water_*/n*_air_ ≈ 1.33 needed to minimize aberrations in remote focusing for an aqueous sample (Supplementary Note 2, Fig. S1). With this lens pair, computational optimization of the oblique-plane imaging geometry yielded center-field tangential and sagittal resolutions of 2.26 *µ*m and 3.08 *µ*m, respectively, across an approximately 5.5 mm field of view (Fig. S2). Guided by this analysis, we designed a monolithic detection module that integrates lens mounting, beam folding, emission filtering and rotational camera positioning into a vertically translatable assembly for focusing along *Z* (Fig. 1B; Supplementary Table 1). We first validated this detection geometry using an externally launched, Powell-lens-based prototype illumination system with resonant beam pivoting for shadow reduction (Supplementary Table 2). When paired with this illumination system, the assembled microscope provides 1.307× magnification over a ~5.4 mm field of view, with measured FWHM resolutions of 2.55 ± 0.19 *µ*m, 3.36 ± 0.19 *µ*m and 17.75 ± 1.4 *µ*m along *X*′, *Y*′ and *Z*′, respectively (Fig. 1F). Having established the performance of the detection architecture, we then designed a compact, baseplate-mounted illumination module to improve mechanical stability and ease of assembly (Fig. 1C–E, Fig. S3, Supplementary Table 3). Experimental characterization of the assembled illumination module yielded a light-sheet thickness of 17.10 *µ*m FWHM along *Z*′ and a lateral extent of approximately 6.5 mm along *X*′ (Fig. S4).

This combination of resolution and field of view was sufficient to resolve subcellular structures in expanded specimens while enabling centimeter-scale expanded tissues to be imaged with as few as three tiles. For example, in 4.2× expanded mouse embryonic fibroblasts labeled for DNA, actin and microtubules, the system resolved individual microtubules (Fig. 1G–I). In chemically cleared, 2-mm-thick sagittal brain slabs from Mobp-eGFP mice, in which oligodendrocytes are fluorescently labeled, the large field of view enabled section-spanning three-dimensional imaging using only two lateral tiles (Fig. 1J). We further imaged an entire mouse liver section labeled for laminin, collagen I and DNA, spanning 5.25 mm before expansion and 21 mm after 4× expansion (Fig. 1K). The orthogonal projection confirmed volumetric coverage through the expanded section (Fig. 1L). The channel-separate zoomed-in views resolved nuclei-dense parenchyma, collagen I-rich stromal features and laminin-positive vascular or basement-membrane structures (Fig. 1M). In a separate region, adjacent collagen I-rich, laminin-poor lumens were consistent with a portal tract containing vascular profiles, such as the portal vein and hepatic artery (Fig. 1N). Together, these examples show that Altair-dvOPM supports imaging across scales, from subcellular detail in expanded cells to centimeter-scale expanded tissues.

Guided by the engineering principles established in Altair-LSFM^12^, Altair-dvOPM brings direct-view oblique-plane imaging into an accessible, open and modular format. By combining photographic-lens-based detection, externally launched oblique illumination and precision-machined baseplates, the platform enables large-field three-dimensional tissue imaging without custom objectives or highly specialized alignment procedures. Complete parts lists, step-by-step online assembly documentation, released CAD files, Zemax models and operation through the open-source microscope-control software *navigate*^13^support replication and extension (Fig. S3; Supplementary Tables 1 and 3). Together, these resources position Altair-dvOPM as a practical platform for three-dimensional mapping of large tissue sections, particularly cleared and expanded specimens.

## Acknowledgements

We would like to acknowledge Grace Gent for their technical assistance in preparing the mouse brain specimens. K.M.D. is supported by the Cancer Prevention and Research Institute of Texas RP250571, the NCI U54CA268072, and the NIGMS RM1GM145399. F.F.V. is funded by a Branco Weiss Fellowship, Society in Science, administered by the ETH Zurich. R.F. is supported by NCI U54CA268072, NIGMS R35GM133522, and NIBIB R01EB035538. S.S.P. is supported by the NINDS R01NS110853. H.Z. is supported by the NIDDK DP1DK139976, and NIAA R01AA028791.

## Author contributions

M.F.: Simulations, CAD design, and Microscope assembly. T.N.: Microscope assembly, Operation, and Analysis. T.D.N: Microscope assembly. J.H.: Simulations, CAD design. Q.S.: Validation. S.G.: Validation. B.C.: Microscope assembly. H.Z.: Validation. S.S.P.: Validation. F.V.: Conceptualization, Methodology, R.F.: Resources, Methodology, Investigation, Writing (review and editing). K.M.D.: Conceptualization, Methodology, Software, Formal Analysis, Writing (original draft, review, and editing), Resources, Visualization, Supervision, Project Administration, Funding Acquisition. All authors were involved in the original draft, review, and editing.

## Data and code availability

Step-by-step assembly documentation for Altair-dvOPM is available at https://thedeanlab.github.io/altair/index.html. CAD files, Zemax models, parts lists and related design files are available at https://github.com/thedeanlab/altair. The microscope control software *navigate* is publicly available on GitHub at https://github.com/TheDeanLab/navigate. The custom software routines used for shearing, registration and image analysis (*clearex*) are publicly available on GitHub at https://github.com/TheDeanLab/clearex. Archived versions of *navigate, clearex* and *altair* will be deposited in Zenodo upon publication. Owing to the size of the imaging datasets, only a representative subset of the data will be archived; the remaining data are available from the corresponding author upon request.

## Supplementary Figures

**Figure S1:**
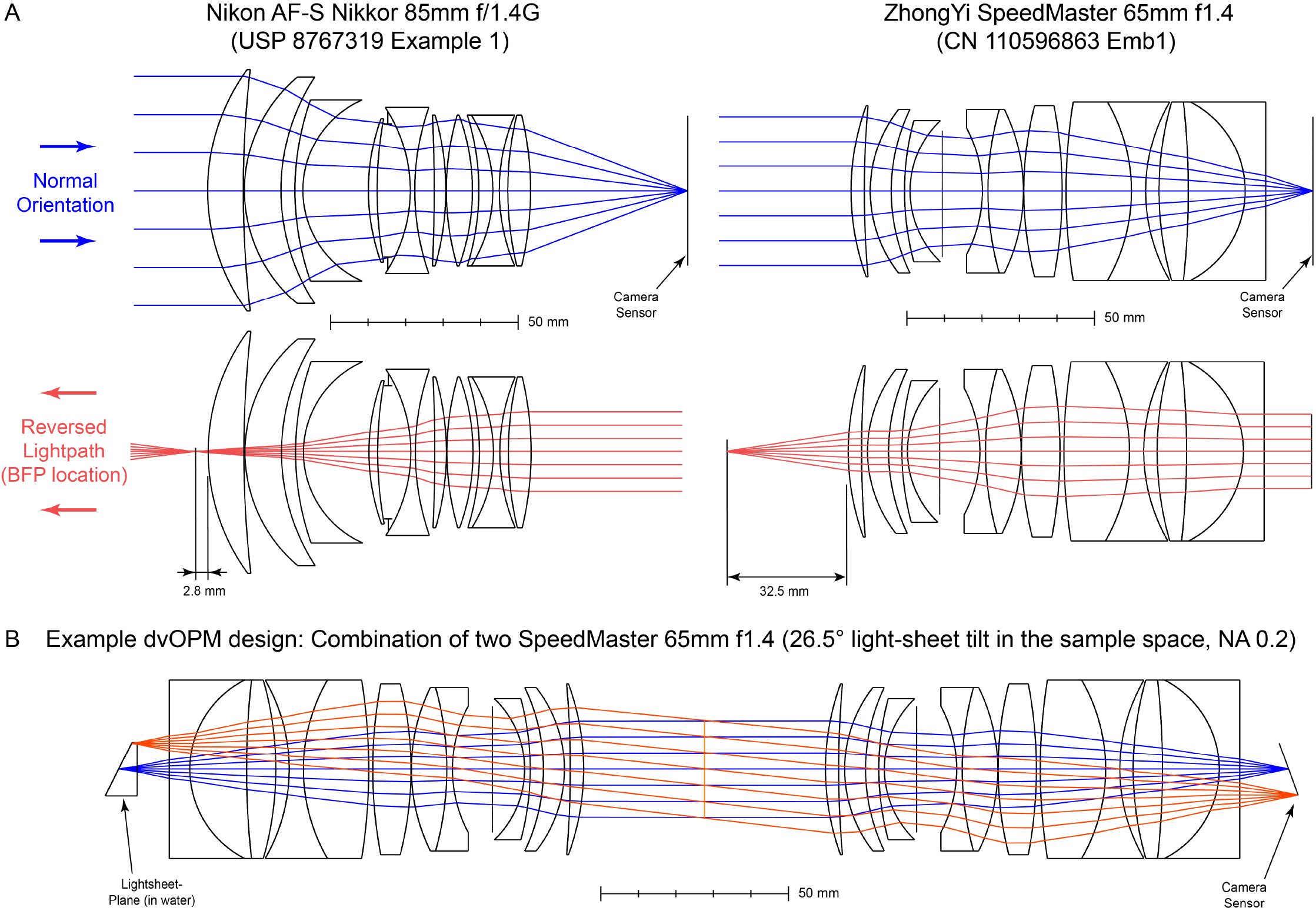
Selection of photographic lenses for Altair-dvOPM. **A**, Location of the nominal image planes in the normal orientation and back-focal-plane positions in the reversed light path for the Nikon AF-S Nikkor 85 mm f/1.4G and ZhongYi Mitakon Speedmaster 65 mm f/1.4 lenses. **B**, Example direct-view OPM design using two ZhongYi Mitakon Speedmaster 65 mm f/1.4 lenses.

**Figure S2:**
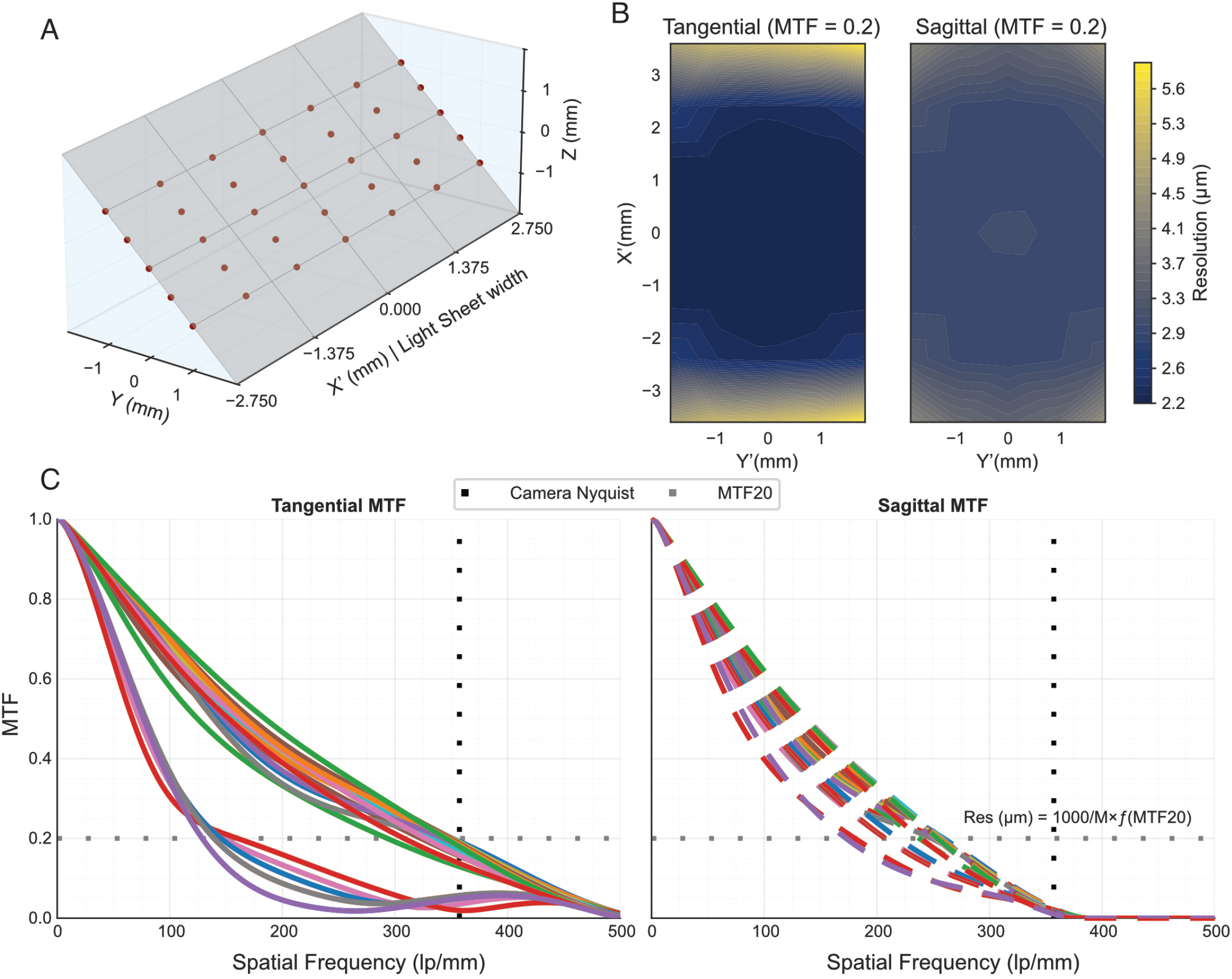
Optical simulations of the detection path. **A**, Oblique image plane used for optimization of the detection optics. Thirty-five field points distributed across *X, Y* and *Z* were assigned weights that decreased with increasing distance from the primary optical axis. These field points correspond to object-space positions that are projected onto the tilted camera sensor. **B**, Simulated resolution maps calculated from the modulation transfer function at 20% contrast (MTF20) in the tangential and sagittal directions. Axes are defined along the oblique image plane shown in **A. C**, Individual tangential and sagittal modulation transfer functions for the field points shown in **A**. Dashed horizontal lines indicate the 20% MTF cutoff used to estimate resolution, and dashed vertical lines indicate the Nyquist sampling limit imposed by the camera pixel size.

**Figure S3:**
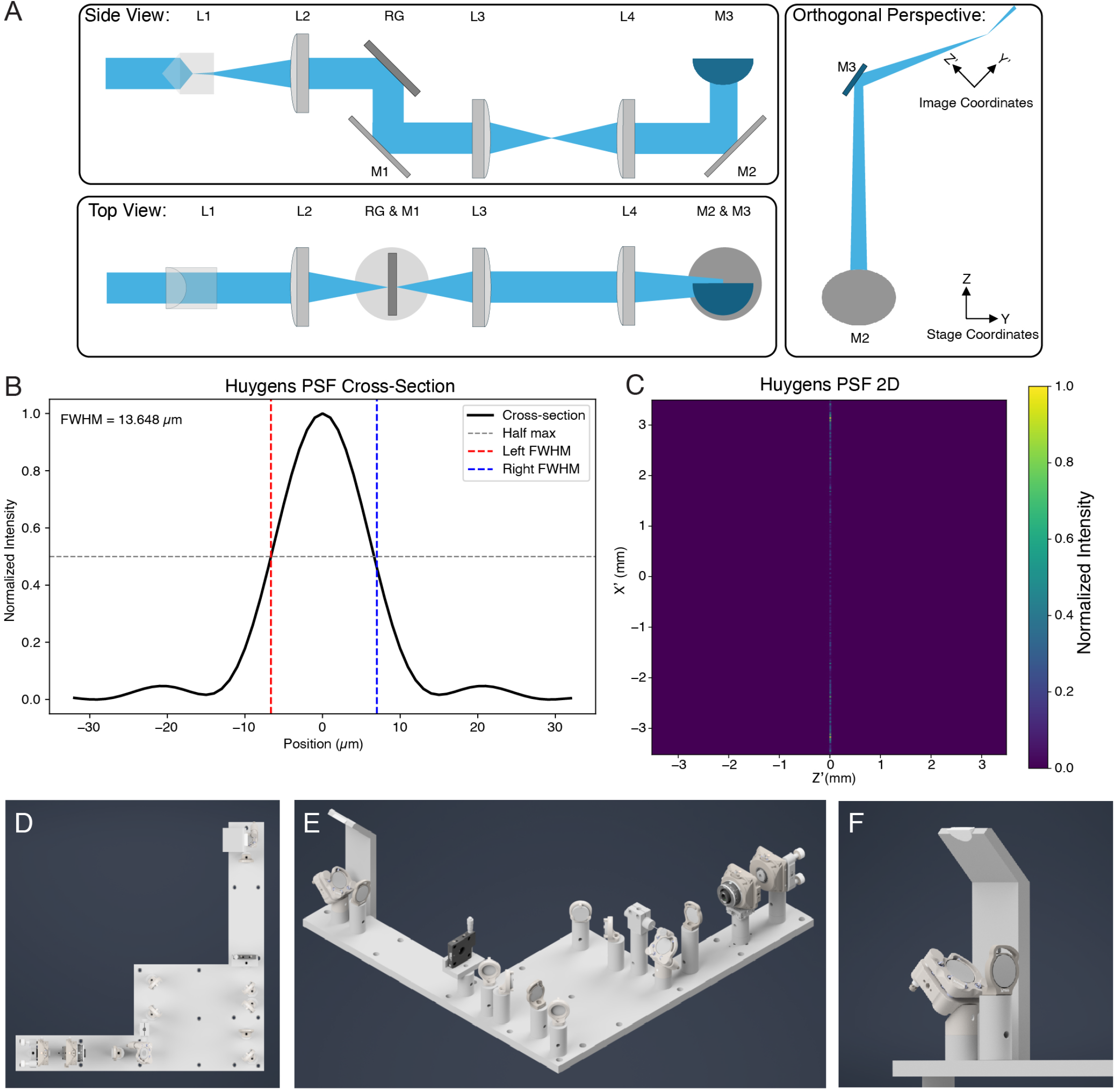
Optical design and simulation of the oblique illumination path. **A**, Optical schematic of the oblique illumination path shown in side view (*Y Z*), top view (*XZ*) and orthogonal perspective (*Y X*). Axes are consistent with those shown in Fig. 1A and are defined in the Zemax coordinate system. A collimated laser beam is focused into a line by a Powell lens (L1), conditioned by an achromatic doublet (L2), reflected from a resonant galvanometer (RG), folded by a mirror (M1) and relayed by achromatic doublets L3 and L4. The beam is then folded vertically by mirror M2 and launched obliquely toward the specimen by mirror M3. **B**, Simulated Huygens point spread function (PSF) cross-section of the illumination beam waist at *λ* = 561 nm. Dashed lines indicate the full width at half maximum (FWHM), corresponding to a predicted light-sheet thickness of 13.6 *µ*m. **C**, Simulated two-dimensional Huygens PSF of the illumination beam along its long, unfocused axis. The illumination beam spans approximately 7 mm. **D**, Overhead view of the custom-machined baseplate for the illumination train. **E**, Isometric view of the illumination baseplate with optical components installed. **F**, Close-up of mirrors M2 and M3. The beam is folded vertically by M2 and then reflected by M3 toward the specimen at the oblique launch angle.

**Figure S4:**
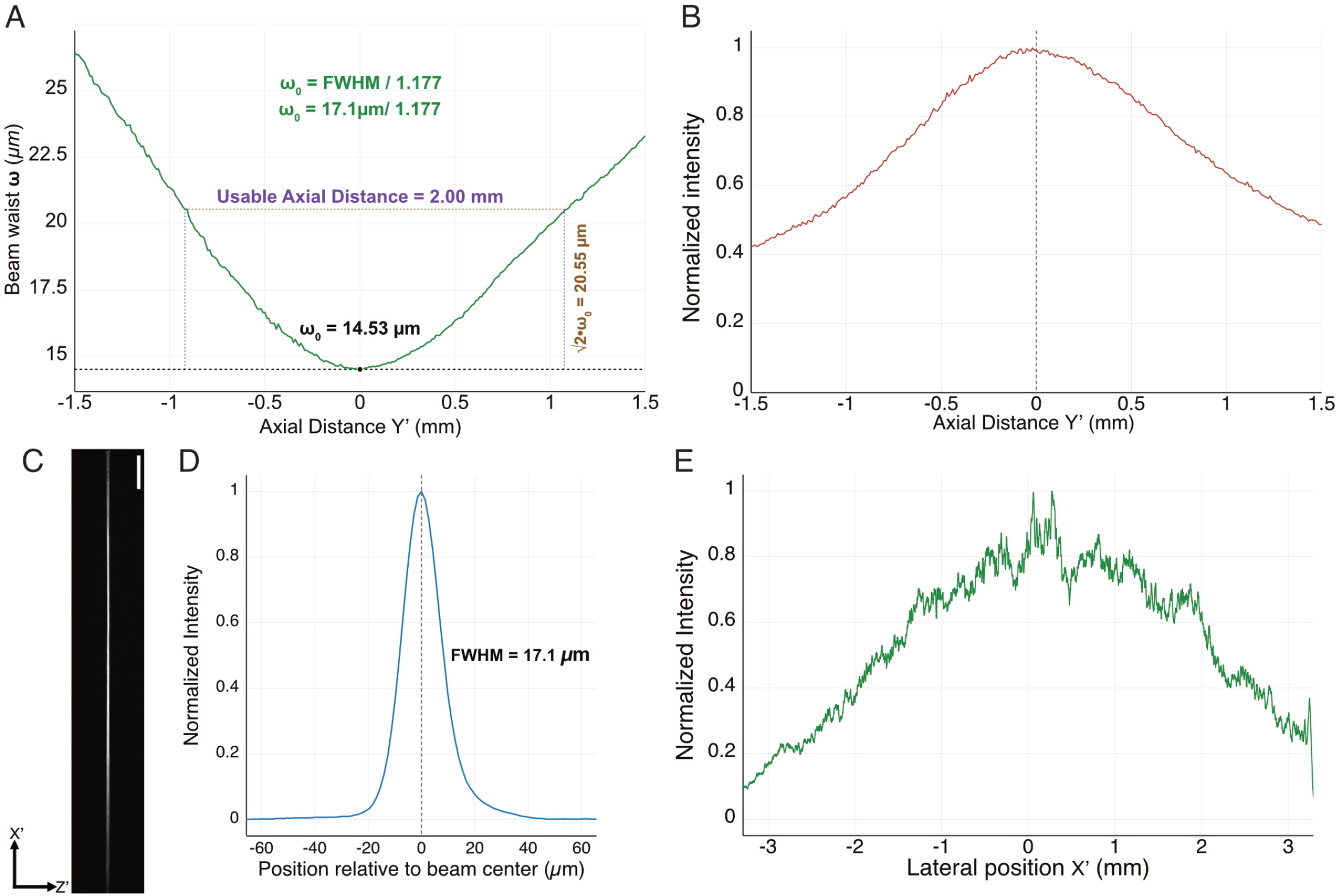
Experimental characterization of the scanned light sheet. **A**, Light-sheet thickness along the propagation axis, *Y*′. The plotted beam waist, *w*, is the Gaussian 1*/e*^2^ intensity radius and was calculated from the measured FWHM using *w* = FWHM*/*1.177. The minimum waist occurred at the best-focus plane, with *w*_0_ = 14.53 *µ*m, corresponding to an FWHM of 17.1 *µ*m. The empirically measured usable axial range, defined as the distance over which the measured waist remained below 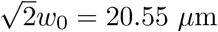, was 2.00 mm. **B**, Normalized peak intensity of the light sheet as a function of propagation distance, *Y*′. The dashed vertical line indicates the focal plane, used as the axial origin. **C**, Camera image of the light sheet at the focal plane. The sheet extends along its long lateral dimension, *X*′, approximately 6.5 mm, and its thickness was measured along *Z*′. Scale bar, 500 *µ*m. **D**, Background-corrected intensity profile along the sheet thickness direction, *Z*′, at the best-focus plane. The measured FWHM was 17.1 *µ*m. **E**, Background-corrected, normalized lateral intensity distribution along the long axis of the light sheet, *X*′, at the best-focus plane. The profile was calculated by summing intensity along *Z*′ for each lateral position, showing the usable lateral extent and relative illumination variation across the sheet.

## Online Methods

### System Design

#### Optical Simulations

Optical simulations for the selection of potential photographic lenses were performed using Ansys Zemax OpticStudio 2023 R1.00 (Ansys, Inc.). Pre-selection of suitable photographic lenses was performed using the online Photons to Photos optical bench tool (Supplementary Note 1). The optical design, modeling, and optimization of the detection and illumination paths were performed in Zemax OpticStudio. In the detection path, the lens files for the photographic lenses were reverse engineered, an object surface representing a 45 ° tilted plane in water, sampled with 35 field points with higher weighting at the central region, with the system aperture defined by an object-space numerical aperture of 0.1786, and simulations performed at a wavelength of 561 nm. Relay geometry and image-plane tilt were optimized in the Zemax Optimization Wizard, using spot size as the optimization criterion and iteratively refining relay distances and camera tilt to minimize aberrations across all weighted field points. Imaging performance was evaluated by calculating the modulation transfer function (MTF) at all field points in tangential and sagittal directions, with resolution defined by the MTF20 criterion in object space. In the illumination path, the Achromatic doublets were modeled using manufacturer-supplied Zemax lens files obtained from the Thorlabs library, and reverse engineered for powell lens. The system was simulated at a wavelength of 561 nm, and the system aperture was defined by pupil diameter of 2 mm. The resonant galvanometer was modeled as a planar mirror. Lenses were added sequentially, with each optimized using the Zemax Optimization Wizard for collimation or focus in its respective dimension before the next element was introduced, building the complete illumination relay. Illumination performance was evaluated in Zemax using Huygens PSF cross-section to quantify light-sheet thickness and 2D Huygens PSF to assess sheet extent and continuity. Simulation files are available on GitHub at www.github.com/DeanLab/altair.

#### CAD and Mechanical Design

The optical system is implemented on two dedicated baseplates, one for detection and one for illumination, with distances and element positions set by the optimized Zemax model. The detection train is built on an L-shaped baseplate in which the filter wheel and right-angle kinematic mirror mount are attached to the vertical panel while the camera stage is anchored to the horizontal plate. This geometry provides a compact, stable path with minimal degrees of freedom in critical alignment axes. Mechanical adaptations in the detection path include a 38 mm to SM2 adapter, a custom camera holder, and a custom camera cap used to replace the Ximea factory cap. The illumination train is built on a primary base slab with an upright plate that terminates in an oblique face for the final beam launch. Most optics are housed in Thorlabs Polaris kinematic mounts with dowel-pin registration for high precision in alignment. Where standard mounts were unsuitable, we designed custom fixtures for the mechanical slit and galvanometer, and custom spacers for the 1XY Powell-lens mount and K1S4 mirror mount. These spacers can alternatively be implemented as custom-height Polaris posts. All custom components were designed in Autodesk Inventor and machined from 6061-T6x aluminum by Xometry, with tolerances of ±0.005 inches. CAD files and ordering information are available on GitHub at www.github.com/DeanLab/altairx.

### Microscope Optics

#### Common microscope hardware

The microscope was built around a shared laser, stage and control architecture used for both prototype and final illumination configurations. Excitation was provided by a LightHub Ultra laser engine equipped with 405 nm LuxX, 488 nm LuxX, 561 nm OPSL and 642 nm LuxX laser lines with nominal powers of 120, 200, 150 and 140 mW, respectively. The laser engine included cleanup filters for the 405, 488 and 642 nm lines, supported TTL-controlled fiber switching and delivered the selected excitation line through high-power FC/APC-terminated fiber outputs with NA = 0.11. Specimens were positioned on an FTP flat-top stage equipped with 4-turn-per-inch drive screws and Heidenhain linear encoders on the *X* and *Y* axes. The lateral stage travel was *<* 130 mm in each axis, with a nominal encoder resolution of 10 nm. Vertical positioning of the stage platform was provided by two parallel LS-50 servo actuators configured for FTP stage-platform focus control, providing a 50 mm travel range with rotary encoder resolution of *<* 10 nm.

#### Illumination System - Initial Prototype

The initial prototype’s illumination path was built using cage components sourced from Thorlabs. Light from the fiber-coupled laser system was collimated with a reflective collimator (Thorlabs, RC08FC-P01.) and directed through a 2 mm pinhole for spatial filtering. The 2 mm pinhole was mounted on a cage-system translation stage (CXY1A, Thorlabs) to enable precise lateral positioning, allowing the beam to be accurately centered on the Powell lens, as it’s highly sensitive to beam size and lateral displacement. The spatially filtered beam was then directed onto a Powell lens with a 10° fan angle (LOCP-8.9R10-2.0, Laserline Optics Canada), generating a uniform line focus that formed a light-sheet with a lateral extent of approximately 7.2 cm at the sample plane. Following the Powell lens, the beam passed through a 30 mm focal length achromatic lens (AC254-030A, Thorlabs) to condition the beam prior to scanning. The illumination was then directed onto a galvanometric mirror (CRS 4 KHz, Novanta), which was used to pivot the light-sheet about the illumination axis. This scanning configuration enabled rapid angular modulation of the excitation direction, similar to multidirectional selective plane illumination microscopy, thereby reducing shadow artifacts arising from absorption and scattering within the specimen. After the galvo, the beam was relayed through a telescope consisting of 100 mm (AC254-100A, Thorlabs) and 125 mm (AC254-125, Thorlabs) focal length lenses to image the pivot point onto the back focal plane of the illumination optics. Finally, the beam passed through a pivoting mount that oriented the illumination at 55° relative to the detection axis, forming an oblique light-sheet suitable for volumetric imaging.

#### Illumination System - Final Version

The final illumination system was redesigned to improve modularity, alignment stability, and compatibility with a compact baseplate architecture. In contrast to the initial prototype, the updated design employs longer focal length optics to accommodate increased propagation distances required for mounting all components on a single baseplate while maintaining consistent optical orientation. This modification facilitates system dissemination and reproducibility, while preserving the beam shaping requirements necessary for light-sheet formation. The following illumination path is visualized in Figure S3A and described in detail below. The excitation source consisted of a LightHub Ultra laser engine coupled into a reflective collimator (CFC11P-A, Thorlabs), producing a collimated multi-line beam. The beam was first directed onto a Powell lens (L1; LOCP-8.9R10-2.0, Laserline Optics Canada) with a 10° fan angle, generating a uniform line profile suitable for light-sheet formation. Following the Powell lens, the beam passed through a 60 mm focal length achromatic doublet (L2; AC254-060A, Thorlabs), which served to condition the beam prior to scanning. The conditioned beam was then incident on a galvanometric mirror (RG; CRS 4 kHz, Novanta), enabling rapid angular pivoting of the light-sheet about the illumination axis. This scanning mechanism provides multidirectional illumination, reducing shadowing artifacts caused by absorption and scattering within the specimen. After reflection from the galvanometric mirror, the beam was directed downward onto a planar mirror (M1; PF10-03-P01, Thorlabs) to restore the optical path orientation. The beam was subsequently relayed through a telescope composed of a 300 mm focal length lens (L3; AC254-300A, Thorlabs) and a 250 mm focal length lens (L4; AC254-250, Thorlabs). This telescope images the pivot point of the galvanometric mirror onto the back focal plane of the illumination optics, ensuring proper angular scanning at the sample plane. A variable mechanical slit (VA100CP, Thorlabs) was positioned within the telescope to control the effective numerical aperture of the illumination, thereby tuning the thickness of the generated light-sheet. To minimize the system footprint while preserving the required optical path length, the beam was folded multiple times within the telescope using planar mirrors (not shown in Figure S3A). Following the telescope, the beam was inverted up using a mirror (M2; PF10-03-P01, Thorlabs) and subsequently reflected by a D-shaped mirror (M3; PFD10-03-P01, Thorlabs). The D-shaped mirror was inclined at 35° relative to the surface normal, resulting in an illumination angle of approximately 20° with respect to the sample stage. This geometry enabled efficient delivery of an oblique light-sheet while maximizing usable space beneath the stage. All optical components were mounted using kinematic mounts with dowel pin registration (Polaris series, Thorlabs) on a custom-designed baseplate. This configuration ensured high mechanical stability, reproducible alignment, and straightforward system assembly, supporting robust operation and ease of dissemination.

#### Detection System

The microscope detection path was built using Mitakon Speedmaster 65 mm f/1.4 and Nikon 85 mm f/1.4 lenses. Although the fast photography lenses have a maximum numerical aperture of 0.25, the effective numerical aperture was reduced to 0.18 to minimize aberrations throughout the field of view. The Mitakon lens (O1) was mounted directly on the filter wheel via a simple adapter stack: 72 mm to 52 mm step-down ring, then SM2A53 to a custom 3D-printed SM2-to-38 mm adapter, which locks into a C60-FW adapter screwed onto the FW1000. The Nikon lens (O2) was coupled to the right-angle kinematic mirror mount through a compact adapter chain: 77 mm to 52 mm step-down ring, followed by a 52 mm to 52 mm adapter and SM2A52, which screws into the SM2-threaded kinematic mount. A 32 mm, 6-position filter wheel (FW1000, Applied Scientific Instrumentation) and a right-angle kinematic mirror mount (KCB2, ThorLabs) with a Ø2” Circular Mirror (PF20-03-P01, ThorLabs) was positioned between the two lenses, contributing a cumulative distance of 136.126 mm between the two lenses. The filter wheel was equipped with Semrock filters for the blue (442/42-32), green (515/30-32), red (595/31-32), and far-red (670/30-32) channels. The detection camera used is MU196MR-ON (Ximea) and it was screwed on a custom machined holder which was mounted on a manual rotation stage (RP005, Thorlabs) placed on a manual linear stage (MT1B, Thorlabs), providing degrees of freedom for fine-tuning the focus and angle in the remote space. The camera’s factory manufactured sensor cap was replaced with custom machined aluminum cap to utilize full field of view when rotated. The camera glass was professionally removed by Pacific Crest Imaging. The full detection assembly was mounted on a custom-manufactured baseplate designed for precise alignment, and vertically positioned using an LS-50 motorized linear stage (Applied Scientific Instrumentation).

## Microscope Control and Data Acquisition

Microscope control and image acquisition are handled by the navigate software package^13^. Stage motion and filter-wheel changes are executed over serial communication using a TigerController (TG8-BASIC, ASI) outfitted with TGCOM, TGFW, and two TGDCM2 control cards. Analog and digital I/O are provided by a PXIe-1073 data-acquisition chassis (NI) with a PXIe-6259 multifunction I/O card and a PXI-6733 analog output card. The system runs on a ProEdge SX6800 workstation (Colfax International) with Windows 10 Pro, an Intel Xeon Silver 4215R CPU at 3.20 GHz, and 96 GB of RAM.

### Sample Preparation

#### Fluorescent Beads

To prepare fluorescent bead samples, we diluted 1 *µ*m YG nanosphere at a concentration of 1:1000 with deionized water and sonicated for 3 minutes to minimize aggregation. The resulting solution was later diluted at a concentration of 1:100 with 2% low-melting agarose for volumetric imaging.

#### Mouse Embryonic Fibroblasts

#### Cell Culture and Fixation

Mouse embryonic fibroblasts (MEFs) were cultured to ~30% confluency on 5 mm glass coverslips pre-cleaned with 70% ethanol. To preserve actin and microtubule architecture, cells were subjected to cytoskeleton-preserving fixation as previously described^14^. Cells were first rinsed with pre-warmed (37 °C) × 1 phosphate-buffered saline (PBS), followed by brief simultaneous permeabilization and fixation using pre-warmed PEM buffer (80 mM PIPES, 5 mM EGTA, 2 mM MgCl_2_, pH 6.8) supplemented with 0.3% Triton X-100 and 0.125% glutaraldehyde for 30 s at 37 °C. Secondary fixation was then performed by replacing the solution with pre-warmed PEM buffer containing 2% paraformaldehyde (PFA) and incubating for 15 min at 37 °C. Following fixation, samples were washed three times with 1 × PBS. Residual aldehydes were quenched with 5 mM glycine in PBS for 10 min at RT. Samples were subsequently stored at 4 °C in 1 × PBS supplemented with 0.02% (w/v) sodium azide (NaN_3_) until further processing.

#### Expansion Microscopy

To physically expand cells, ultrastructure expansion microscopy was performed^15^. Fixed cells were rinsed three times with 1 × PBS to remove NaN_3_ and incubated in an anchoring solution containing 1.4% formaldehyde (FA, Sigma-Aldrich) and 2% acrylamide (AA, Bio-Rad) in 1 × PBS for 3 h at 37 °C without agitation. Following anchoring, samples were incubated in monomer solution consisting of 23% sodium acrylate (SA, AmBeed), 10% AA, 0.1% N,N′-methylenbisacrylamide (BIS, Sigma-Aldrich), 0.5% tetramethylethylenediamine (TEMED, Bio-Rad), 1 × PBS and Milli-Q water for 30 min on ice to allow uniform monomer diffusion. Coverslips were then inverted (cell-side down) onto an ice-cooled gelation chamber containing ~15 *µ*L of monomer solution supplemented with 0.5% ammonium persulfate (APS). After initial polymerization on ice, the samples were transferred to 37 °C and incubated for 45 min under humidified conditions for complete gel formation. To homogenize cellular ultrastructure and enable isotropic expansion, hydrogels were incubated in denaturation buffer (200 mM SDS, 200 mM NaCl, 50 mM Tris-Base, and Mili-Q water, pH 9) at 95 °C for 1.5 h. Gels were subsequently transferred to excess Milli-Q water and allowed to expand overnight at RT to ensure complete removal of denaturation agents and full physical enlargement.

#### Immunofluorescence Labeling

Expanded gels were blocked at RT for 1 h in 3% bovine serum albumin (BSA) and 0.01% Triton X-100 in 1 × PBS. For indirect immunofluorescence, samples were incubated for 2.5 h at 37 °C with primary antibodies diluted in staining buffer (1% BSA + 0.01% Triton X-100 in 1 × PBS): rat anti-*α*-Tubulin (Selleckchem, #F1566, 1:200) and mouse anti-*β*-Actin (Sigma-Aldrich, #A1978, 1:200, RRID: AB_476692) with gentle agitations. After three washes with PBST (1 × PBS containing 0.01% Triton-X, 20 min each), cells were incubated for 2 h with secondary antibodies diluted in staining buffer: donkey anti-rat Alexa Fluor 488 (Invitrogen, #A-21208, 1:500, RRID: AB_2535794) and donkey anti-mouse CF555 (Biotium, #20037, 1:500, RRID: AB_1085438). For nuclear staining, DAPI (0.3 *µ*M, Thermo Fisher Scientific, #62248) was applied for 15 min at RT.

### Mouse Specimens

#### Vertebrate Animals

Animal work described in this manuscript was approved by and conducted under the oversight of the UT Southwestern Institutional Animal Care and Use Committee. TetO-Cas9/rtTA mice on a C57BL/6J background were purchased from The Jackson Laboratory (strain 029415). Mobp-eGFP mice^16^were purchased from the Mutant Mouse Resource & Research Centers. All mice were maintained at UT Southwestern Medical Center in a temperature-, humidity-, and light-controlled animal facility with access to food and water ad libitum. Mice were transcardially perfused with 0.01 M phosphate-buffered saline followed by 4% paraformaldehyde in 0.1 M phosphate buffer. Brains were postfixed in 4% paraformaldehyde overnight at 4 °C and then washed repeatedly with PBS containing 0.02% sodium azide prior to clearing.

#### Liver Specimen

Liver samples were obtained from C57BL/6J mice. Animals were euthanized, and livers were immediately excised and fixed in 4% paraformaldehyde (Thermo Fisher Scientific, #AJ19943K2) overnight at 4 °C with gentle agitation. Fixed tissues were embedded in 2% agarose and sectioned into 100 *µ*m slices. For antigen retrieval, sections were subjected to heat-induced epitope retrieval (HIER) in Tris–EDTA buffer (Enzo Life Sciences, #ENZ-ACC113) at 95 °C for 30 min, followed by three washes in PBS (30 min each). Samples were then blocked for 2 h at room temperature in PBST containing 4% bovine serum albumin (BSA; Equitech-Bio, BAH65). Primary antibodies were diluted in blocking buffer and applied overnight at 4 °C with gentle agitation: anti-laminin (LAMA1) (Sigma-Aldrich, L9393, RRID: AB_477163, 1:100) and anti-collagen I (Invitrogen, MA1-26771, RRID: AB_2081889, 1:100). After incubation, sections were washed three times in PBST (1 h each). Fluorophore-conjugated secondary antibodies, goat anti-rabbit IgG (Alexa Fluor 568; Thermo Fisher, #A-11036) and goat anti-mouse IgG (Alexa Fluor 488; Thermo Fisher, #A-11029), were applied at 1:100 dilution for 12 h at 4 °C, followed by three additional PBST washes (1 h each). Sections were anchored overnight at room temperature in 0.1 mg mL^*−*1^Acryloyl-X, SE (Thermo Fisher Scientific) in PBS. Samples were then washed twice in PBS for 15 min each and incubated overnight at 4°C in monomer solution containing 19% (w/w) sodium acrylate, 10% (w/w) acrylamide and 0.1% (w/w) *N, N*′-methylenebisacrylamide. For gelation, samples were gently transferred into a gelation chamber composed of a microscope slide and silicone spacer (McMaster-Carr, #6459N112), and overlaid with gelation solution consisting of monomer solution supplemented with 0.5% (w/w) ammonium persulfate and 0.5% (w/w) tetramethylethylenediamine. Gelation proceeded on ice for 30 min, followed by 1.5 h at 37°C in a humidified incubator. Polymerized samples were denatured for 6 h at 37°C in digestion buffer containing 50 mM Tris, pH 8.0, 1 mM EDTA, 0.5% Triton X-100 and 1 M NaCl, supplemented with 8 U mL^*−*1^proteinase K (MilliporeSigma, #39450-01-6). After three 30-min washes in PBS, nuclei were stained with SYTOX™ Green (Thermo Fisher, #S7020; 1:1000 in PBS) for 3 h at room temperature. Samples were then washed three additional times in PBS for 1 h each. Before imaging, samples were expanded in deionized water for at least 1 h, with three exchanges of fresh deionized water. The estimated linear expansion factor was approximately 4×.

#### Brain Specimen

Brain samples were obtained from Mobp-GFP mice, which express eGFP in oligodendrocytes. Brain samples were pretreated with 50% CUBIC-L (product no. T3740, TCI Chemicals) with gentle rotation at 37 °C for 24 h. The samples were then incubated in 100% CUBIC-L with gentle rotation at 37 °C for 72 h for delipidation, with the CUBIC-L refreshed every 24 h. After delipidation, the brains were washed three times with 1× PBS, with the buffer refreshed every 2 h at RT. For immunolabeling, brain samples were permeabilized overnight at RT with gentle rotation in blocking buffer (0.5% NP-40, 10% DMSO, 5% donkey serum, and 0.5% Triton X-100 in 1× PBS). The endogenous GFP signal was then amplified by incubation with anti-GFP antibody (AB_2307313) for 6 days at RT with gentle rotation. The samples were washed with wash buffer (0.5% NP-40 and 10% DMSO in 1× PBS) for 6 h, with the buffer refreshed every 2 h, and were then post-fixed in 4% PFA overnight at RT to reduce fluorescent aggregate formation. The brains were then labeled with secondary antibody (donkey anti-chicken Alexa Fluor 488, cat. no. A78948) for 3 days at RT with gentle rotation. Following secondary labeling, the samples were washed with wash buffer for 6 h, with the buffer refreshed every 2 h. After immunolabeling, the mouse brains were pretreated with 50% CUBIC-R+ (product no. T3741, TCI Chemicals) for 1 day at RT with gentle rotation. The next day, the brain tissue was incubated in 100% CUBIC-R+ to match the refractive index.

## Data Analysis

### Image Post-Processing

Raw image datasets were processed in ClearEx after materialization to a canonical OME-Zarr store. Illumination nonuniformity was corrected independently for each position and channel using a tiled flat-field model with 256×256-pixel fitting tiles, lateral halo blending (32 pixels in x and y), smoothness parameter 1.0, and working size 128, without dark-field estimation. The corrected volumes were then deskewed into orthogonal coordinates by applying a 45°yz shear with automatic rotation, linear interpolation, zero-value padding, and 2-voxel ROI padding, yielding rectified volumes at 5.0 ×1.0599×1.0599 *µ*m voxel spacing. For multiposition datasets, the rectified tiles were registered at full resolution using a designated reference channel and a similarity transform model, with FFT-based initialization followed by multiresolution ANTs optimization (200/100/50/25 iterations; sampling rate 0.2) over 128-voxel pairwise overlap regions. Registered tiles were fused in 256^3^-voxel chunks with gain-compensated feather blending across 400-voxel overlap bands (blend exponent 0.5; gain clipping 0.5–2.0) to generate the final stitched volume. For single-position datasets, the rectified volume was carried forward without registration or stitching. For quality control and figure generation, orthogonal maximum-intensity projections (xy, xz, yz) were exported as OME-TIFF files from the rectified or stitched volumes, with z resampled to the lateral pixel size for the xz and yz views. ClearEx is available as an open-source package at www.github.com/DeanLab/ClearEx.

### Resolution Analysis

To analyze the resolution of the system, we imaged 1 *µ*m green fluorescence beads embedded in agarose on a coverslip-bottom petri dish. Volumetric stacks were acquired by scanning the agarose volume through the illumination light-sheet over a 400 *µ*m range at 2 *µ*m step size. Image stacks were deskewed using BigSticher in ImageJ and its PSF over entire 3D volume was evaluated with our own code in Matlab. Individual beads were first detected using intensity thresholding and centroid localization within the 3D volume. For each detected bead, sub-volumes centered on the bead were extracted and analyzed to estimate the PSF width along the lateral (x–y) and axial (z) directions. The full width at half maximum (FWHM) of the intensity profiles was calculated by fitting the bead intensity distribution with a Gaussian model along each axis. Measurements from all valid beads were aggregated, and the median resolution values were reported along with the corresponding standard deviation. To evaluate the lateral uniformity of the imaging performance, beads located at different positions across the field of view were analyzed. Resolution measurements were mapped as a function of lateral position to assess potential field-dependent aberrations or performance degradation toward the edges of the imaging field.

## Supplementary Tables

**Table 1.** Detection Train Parts List.

| Part Number | Source | Qty | Cost | Description |
| --- | --- | --- | --- | --- |
| **Optics** |  |  |  |  |
| Mitakon Speedmaster 65mm f/1.4 | Mitakon | 1 | \$599.00 | Fast photographic lens |
| AF-S NIKKOR 85mm f/1.4G | Nikon | 1 | \$1159.95 | Fast photographic lens |
| MU196MR-ON | Ximea | 1 | \$900.00 | Monochrome camera, AR2020 sensor |
| FW-1000 | ASI | 1 | \$3700.00 | High-speed motorized filter wheel |
| PF20-03-G01 | Thorlabs | 1 | \$115.68 | Ø2" protected aluminum mirror |
| **Optomechanics** |  |  |  |  |
| C60-FW | ASI | 2 | \$200.00 | Filter wheel adapter |
| KCB2 | | 1 | \$204.38 | Kinematic mirror mount |
| SM2A52 | Thorlabs | 1 | \$25.29 | SM2 to M52 adapter |
| SM2A53 | Thorlabs | 1 | \$25.29 | M52 to SM2 adapter |
| 77mm to 52mm lens adapter | Generic | 1 | \$9.99 | 77mm to 52mm step-down ring |
| 72mm to 52mm lens adapter | Generic | 1 | \$6.79 | 72mm to 52mm step-down ring |
| 52mm to 52mm lens adapter | Generic | 1 | \$5.95 | 52mm male-to-male coupler |
| MSAP90 | Thorlabs | 1 | \$55.83 | Right-angle plate |
| **Stages** |  |  |  |  |
| S561-2235B | ASI | 1 | \$8200.00 | Flat-top stage |
| Dual-LS-50-FTP | ASI | 1 | \$5100.00 | Two parallel actuators with LS-5007 |
| LS-5008-K | ASI | 2 | \$700.00 | 90° brackets for Dual-LS-50-FTP |
| LS-50-AMERL | ASI | 1 | \$2550.00 | Motorized linear stage |
| MT1B | Thorlabs | 1 | \$278.36 | Manual linear stage |
| RP005 | Thorlabs | 1 | \$99.69 | Manual rotation stage |
| PT1B | Thorlabs | 2 | \$268.20 | Manual linear stage |
| **Modular Baseplate & Custom 3D Printing** |  |  |  |  |
| Baseplate | Xometry | 1 | \$300.00 | L-Shaped main baseplate |
| Custom Adapter | Xometry | 1 | \$100.00 | 38mm to SM2 adapter |
| Custom Camera Holder | Xometry | 1 | \$200.00 | Holder for camera |
| Custom Camera Cap | Xometry | 1 | \$200.00 | Redesigned camera cap |

**Table 2.**
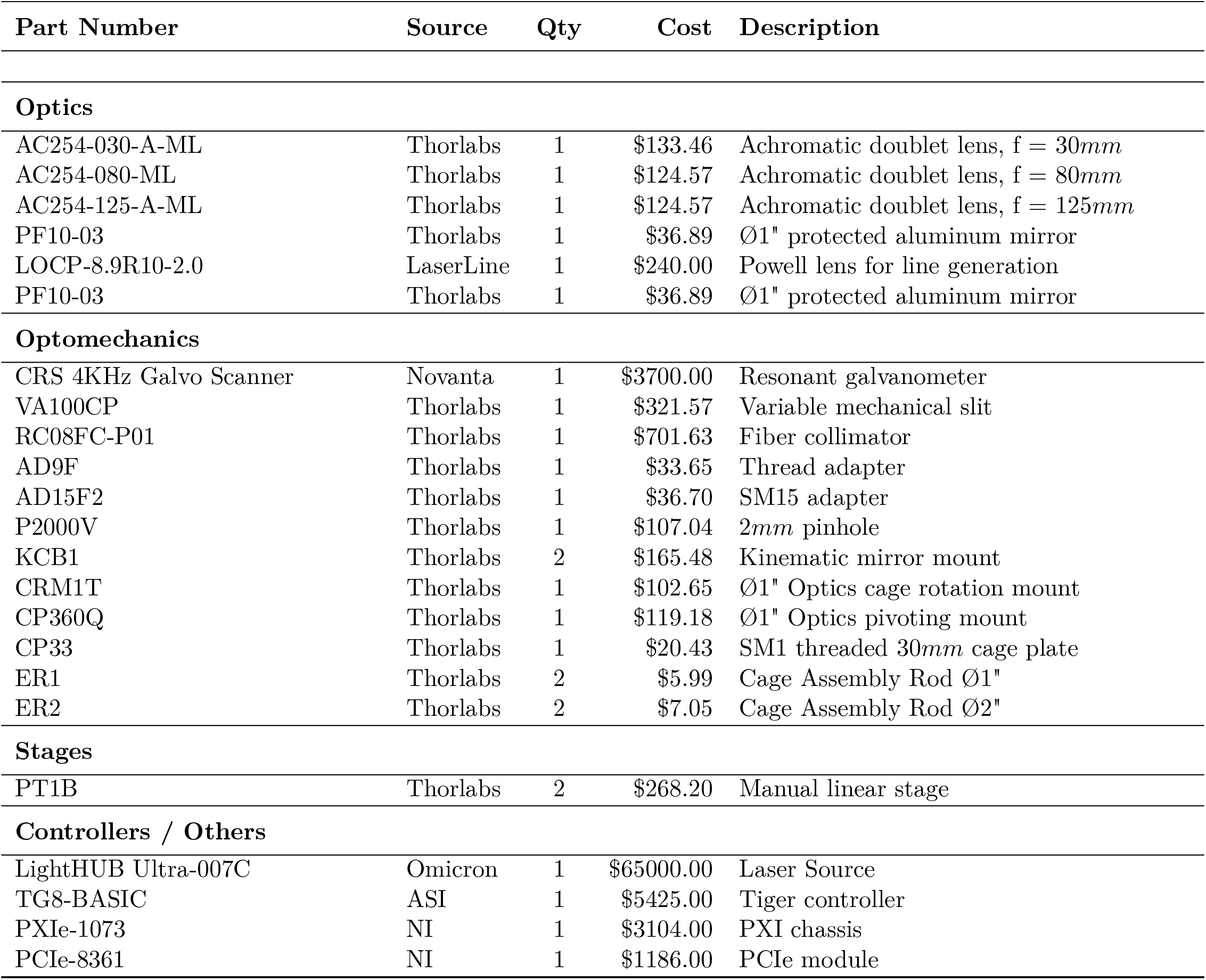
Illumination Train Parts List (Initial Prototype)

**Table 3.** Illumination Train Parts List (Final Prototype)

| Part Number | Source | Qty | Cost | Description |
| --- | --- | --- | --- | --- |
| **Optics** |  |  |  |  |
| AC254-060-A | Thorlabs | 1 | \$95.34 | Achromatic doublet lens, $f = 60mm$ |
| AC254-250-A | Thorlabs | 1 | \$95.34 | Achromatic doublet lens, $f = 250mm$ |
| AC254-300-A | Thorlabs | 1 | \$95.34 | Achromatic doublet lens, $f = 300mm$ |
| PF10-03 | Thorlabs | 7 | \$36.89 | Ø1" protected aluminum mirror |
| LOCP-8.9R10-2.0 | LaserLine | 1 | \$240.00 | Powell lens for line generation |
| PFD10-03-P01 | Thorlabs | 1 | \$71.50 | Ø1" protected silver D-shaped mirror |
| **Optomechanics** |  |  |  |  |
| PLS-P150 | Thorlabs | 2 | \$39.10 | Polaris post, 1.5" |
| PLS-P2 | Thorlabs | 5 | \$42.00 | Polaris post, 2" |
| PLS-P3 | Thorlabs | 3 | \$46.12 | Polaris post, 3" |
| POLARIS-K1XY | Thorlabs | 1 | \$1473.39 | XY kinematic mount |
| POLARIS-1XY | Thorlabs | 3 | \$1088.67 | Compact XY mount |
| POLARIS-B1F | Thorlabs | 5 | \$170.26 | Fixed mount base |
| POLARIS-B1S | Thorlabs | 3 | \$119.50 | Slim mount base |
| POLARIS-MA45 | Thorlabs | 2 | \$53.69 | 45° adapter |
| POLARIS-K1S4 | Thorlabs | 2 | \$198.35 | Kinematic mount for 1" optics |
| CRS 4KHz Galvo Scanner | Novanta | 1 | \$3700.00 | Resonant galvanometer |
| VA100CP | Thorlabs | 1 | \$321.57 | Variable mechanical slit |
| CFC11P-A | Thorlabs | 1 | \$338.29 | Fiber collimator |
| LRM1 | Thorlabs | 1 | \$109.65 | Lens adapter |
| AD9F | Thorlabs | 1 | \$33.65 | Thread adapter |
| AD15F2 | Thorlabs | 1 | \$36.70 | SM15 adapter |
| P2000K1 | Thorlabs | 1 | \$87.74 | 2mm pinhole |
| RS2.5P | Thorlabs | 8 | \$35.91 | 2.5" pedestal post |
| **Stages** |  |  |  |  |
| XP10X | Thorlabs | 1 | \$620.00 | Manual linear stage |
| XP10-C1 | Thorlabs | 1 | \$90.00 | Actuator for XP10X |
| LNR25M | Thorlabs | 1 | \$828.75 | Manual linear stage |
| **Controllers / Others** |  |  |  |  |
| LightHUB Ultra-007C | Omicron | 1 | \$65000.00 | Laser Source |
| TG8-BASIC | ASI | 1 | \$5425.00 | Tiger controller |
| PXIe-1073 | NI | 1 | \$3104.00 | PXI chassis |
| PCIe-8361 | NI | 1 | \$1186.00 | PCIe module |
| **Modular Baseplate & Custom 3D Printing** |  |  |  |  |
| Baseplate | Xometry | 1 | \$300.00 | L-Shaped main baseplate |
| Side Plate | Xometry | 1 | \$100.00 | Attachment to the main baseplate |
| Custom Holder for Galvo | Xometry | 1 | \$100.00 | Holder for Galvanometer |
| Custom Holder for Slit | Xometry | 1 | \$100.00 | Holder for Mechanical Slit |
| Custom Spacer (K1S4) | Xometry | 2 | \$50.00 | Spacer Kinematic Mount |
| Custom Spacer (1XY) | Xometry | 1 | \$50.00 | Spacer 1XY Mount |

### Supplementary Note 1 – Principles of Macroscale Oblique Plane Microscopy

Oblique plane microscopy (OPM) and related swept implementations such as SCAPE comprise a family of light-sheet-based imaging systems in which fluorescence from an obliquely illuminated plane is detected through a specimen-interfacing objective^1,2^. In canonical single-objective OPM and SCAPE implementations, the oblique illumination is generated through the same objective that collects fluorescence. In related mesoscopic and direct-view architectures, the illumination may instead be launched externally or through a non-orthogonal illumination objective, while detection still proceeds by imaging an oblique plane through the primary detection objective^5–7,11^. The resulting tilted image plane is transferred through a remote-refocusing system for camera-based detection. This geometry preserves the optical sectioning and volumetric throughput advantages of light-sheet microscopy while maintaining epi-style or open-top sample access and compatibility with conventional sample formats. Here, we focus on mesoscopic or macroscale OPM implementations with usable fields of view of approximately 1 mm or larger, as these systems are most relevant to large cleared and expanded specimens.

The conceptual basis for OPM was established by Dunsby, who introduced a remote-refocusing strategy for imaging an oblique plane within the specimen onto a planar detector^1^. Bouchard *et al*. subsequently demonstrated SCAPE, showing that the oblique sheet could be swept through the specimen using a stationary specimen-facing objective to enable high-speed volumetric imaging^2^. These studies established the practical importance of single-objective light-sheet imaging, but scaling the approach to mesoscopic fields of view introduces a central optical constraint. In conventional remote-refocused OPM, fluorescence from the oblique intermediate image plane is reimaged by a tilted tertiary objective. Because objective design imposes an inverse relationship between numerical aperture and field of view, large-format imaging favors low-NA reimaging optics. However, low-NA tertiary objectives inefficiently collect the obliquely propagating fluorescence, creating a tradeoff between field of view, collection efficiency and axial resolution^6,9^.

Several architectures have addressed this limitation by modifying the remote-refocusing or reimaging geometry. SCAPE 2.0 improved optical throughput and expanded the corrected image circle, enabling millimeter-scale volumetric imaging while retaining micrometer-scale sampling and high volumetric imaging speed^3^. DaXi used a custom remote-focusing objective and multiview acquisition to extend single-objective light-sheet microscopy to larger imaging volumes while maintaining submicrometer lateral and low-micrometer axial resolution^17^. Hoffmann and Judkewitz introduced diffractive OPM, in which a blazed diffraction grating is placed at the intermediate image plane to redirect fluorescence into the acceptance cone of a low-NA tertiary imaging system^9^. This approach enabled OPM with a 0.28 NA objective over a 3.3 × 3.0 × 1.0 mm^3^addressable volume, although at the cost of reduced photon efficiency and axial resolution^9^. A later blazed OPM implementation extended this diffractive-reimaging concept to brain-wide neuronal activity imaging in an adult vertebrate^10^. Importantly, these diffractive approaches modify the direction and collection efficiency of the oblique intermediate image, but still rely on a downstream imaging system to form the camera image.

A complementary strategy is to modify the illumination geometry while preserving oblique-plane detection. Shao *et al*. introduced a mesoscopic OPM with a diffractive light sheet, achieving fields of view up to 5.4 × 3.3 mm with approximately 2.5 × 3 × 6 *µ*m resolution and enabling large-scale volumetric imaging of zebrafish larvae^18^. Other approaches decouple illumination and detection more explicitly by using a second, non-orthogonal illumination objective or sample-side optical element. Singh *et al*. described an open-top, non-orthogonal dual-objective OPM for freely moving organisms, demonstrating millimeter-scale volumetric imaging with cellular-level resolution^5^. Daetwyler *et al*. used a microprism to reflect the illumination sheet into the specimen in combination with a transmission grating to redirect the fluorescence into a tilted tertiary imaging system, enabling mesoscopic OPM with improved sectioning uniformity and reduced shadowing across larger volumes^7^. Davis *et al*. subsequently combined a non-orthogonal dual-objective OPM geometry with axial sweeping of the illumination waist, improving axial confinement and resolution uniformity over a mesoscopic field of view^6^. In this implementation, synchronization of an electrically tunable lens with the rolling shutter of the camera enabled near-isotropic imaging of freely moving *Nematostella* over a 1.0 × 0.7 × 0.4 mm volume at 1.7 × 2.6 × 3.7 *µ*m resolution and 0.5 Hz volume rate^6^.

Direct-view OPM addressed the tertiary-objective bottleneck more directly by eliminating the tertiary imaging system and placing the camera sensor in the remote image space^11^. This arrangement avoids the low-NA collection penalty associated with tilted tertiary reimaging and enables efficient detection at low numerical aperture. Using a commercially available camera with small pixels, Lamb *et al*. demonstrated imaging over a 5.3 × 3.0 × 2.6 mm^3^volume with approximately 2 *µ*m lateral and 22 *µ*m axial resolution^11^. However, this design still relies on a pupil-mapped relay between the primary and secondary objectives to satisfy the requirements of remote refocusing. Thus, although direct-view OPM removes the tertiary objective, it does not eliminate the alignment and pupil-conjugation constraints of the conventional remote-refocusing train.

Together, these developments show that mesoscopic OPM has advanced along several complementary axes: improved objective-coupled remote refocusing, diffractive or direct-view reimaging, engineered illumination and non-orthogonal open-top geometries. The present work builds on this literature by using externally accessible pupils in fast photographic lenses to construct a compact direct-view detection path, while retaining externally launched oblique illumination suitable for large cleared and expanded specimens.

## Supplementary Note 2 – Survey of Fast Photographic Lenses with Externalized Pupils

For the design of our direct-view OPM (dvOPM), we identified a variety of conditions for the selection of possible lenses:

- We opted for a combination of two objectives (O1 and O2 – essentially an infinity objective & tube lens combo as in an infinity-corrected microscope) to have a parallel lightpath inbetween O1 and O2. This allows for the integration of optical filters and dichroics with coatings optimized for parallel light and flexible selection of the total magnification of the system.
- The total magnification *M* of the system should be 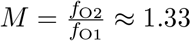 to fulfil the condition for aberration-free remote focusing and relaying the image from water to a camera in air.
- In order to have optimal coupling of emitted light from O1 to O2, both objectives should have an external pupil/stop location/back focal plane. This means that the exit pupil of O1 can be superimposed with the entrance pupil of O2. This would also allow placing a scan mirror inbetween O1 and O2 for fast scanning (if necessary).
- The sample-sided working distance of the objective O1 has to be sufficiently high to accommodate typical sample holders and stages.
- The image-sided working distance of the objective O2 has to be sufficiently high to accommodate a tilted camera and its housing.

The NA/f-number of the objectives should be as high as possible (ideally better than f/2.5 ≈ NA 0.2) for the best possible resolution and light-collection efficiency.

1. The whole system makes use of the Scheimpflug principle to image a tilted lightsheet onto a tilted camera chip.
2. The FOV of the system should be larger than 4 mm, ideally as large as possible.

Unlike scan lenses, photographic objectives are usually not designed with an external aperture stop. Nonetheless, many commercial objectives have mechanically accessible stop locations / back focal plane (BFP) positions. To screen for possible combinations, we used an online optical bench tool that has a large underlying lens database assembled from patent instructed. By toggling the “MarginalI” and “MarginalO” buttons, it is possible to visualize the location of the entrance and exit pupils.

As a first step, we screened for possible O1 objectives with focal lengths of 50–100 mm to minimize the mechanical size of the system. We only considered objectives with fixed focal lengths (no zoom objectives) as zoom lenses have pupil locations that change location depending on the zoom position. One option is the Mitakon Speedmaster Zhong Yi 65mm F1.4 GFX: It is designed for Fujifilm GFX medium-format cameras with an image format of 43.8 × 32.9 mm – much larger than our desired FOV (Optical Bench Link). In addition, it is designed with a flat surface towards the sensor (or towards the sample in our application) which makes cleaning of this surface (e.g., due to spilled immersion fluid) and tip/tilt alignment relative to the sample holder much easier. Its BFP/entrance pupil is located 32.5 mm away from the front lens. We reoptimized the lens in using Ansys Zemax OpticStudio 2023 R1.00 (Ansys, Inc.) based on the patent information and simulated its performance in a symmetric dvOPM configuration with M=1.0 (Fig. S1). Under these circumstances, the system was capable of achieving a NA of 0.20 with imaging quality close to the diffraction limit (max. RMS wavefront error of 0.14*λ* at 589 nm up for image locations up 7 mm off-axis, highlighting its suitability for large-FOV microscopy applications.

To achieve a magnification *M* ≈ 1.33, we decided to pair this objective with an 85 mm objective to reach *M* = 85mm*/*65mm = 1.307. A suitable option is the Nikon 85 mm f1.4G (USP8767319) single-lens reflex (SLR) objective (Optical Bench Link). Its BFP is located 2.8 mm in front of the first element. However, one can increase the distance between the objectives from (32.5 + 2.8 = 35.3 mm) if one accepts that the optical path is not telecentric on the camera side. We made use of this property to add a 2” fold mirror and filter wheel into the optical path between O1 and O2.

